# The speed-accuracy tradeoff reveals flexible access to accumulating sensory evidence during human decision making

**DOI:** 10.1101/420430

**Authors:** Stephanie Nelli, Sirawaj Itthipuripat, Nuttida Rungratsameetaweemana, John T. Serences

## Abstract

Decisions made about identical perceptual stimuli can be radically different under changing task demands. For example, the need to make a fast decision undermines the accuracy of that decision, a well-documented effect termed the speed-accuracy tradeoff (SAT). Models of the SAT are generally based on theories of decision making in which responses are triggered only after sensory evidence accumulation terminates at a set threshold. Within this accumulate-to-bound framework, speed pressure operates by lowering the response threshold, allowing for faster responses at the expense of accumulated sensory evidence. To empirically examine the mechanisms necessary for adaptively adjusting the speed and accuracy of decisions, we used an event-related potential that indexes sensory evidence accumulation in the human brain. Instead of lowering response thresholds, we found that speed pressure adaptively shifts responses to occur close to where the rate of evidence accumulation peaks. Moreover, responses are not triggered automatically by the termination of the accumulation process, as sensory evidence continues to build after speeded decisions. Together these results suggest that response processes adaptively access accumulating sensory evidence depending on task demands and support parallel over serial models of decision making.

## Introduction

Current task demands often prioritize either the speed or accuracy of a decision at the expense of the other (the speed-accuracy tradeoff, or SAT) (Heitz, 2014; Heitz & Schall, 2012; Henmon, 1911; Roger Ratcliff & Rouder, 1998; Shadlen, Hanks, Churchland, Kiani, & Yang, 2006). For example, classifying whether a vehicle on the highway is law enforcement might inspire a fast decision to slow down. While this speeded classification is less likely to be accurate, it is nevertheless a good strategy for avoiding a costly ticket.

The SAT is typically modeled within the serial accumulate-to-threshold framework of decision making, which holds that responses are only executed once sensory information accumulates to a specific threshold (Bogacz, Brown, Moehlis, Holmes, & Cohen, 2006; Bogacz, Wagenmakers, Forstmann, & Nieuwenhuis, 2010; Brody & Hanks, 2016; R. Kiani, Hanks, & Shadlen, 2008; Roger Ratcliff, Smith, Brown, & McKoon, 2016). In these models, speeded decisions are due to lower response thresholds, meaning decisions are based on impoverished sensory evidence and are therefore less accurate (Drugowitsch, Moreno-Bote, Churchland, Shadlen, & Pouget, 2012). In contrast, increased response thresholds allow for the accumulation of more sensory evidence and thus slower but more accurate decisions (Heitz, 2014; Shadlen et al., 2006).

Several neural markers have been used to index the accumulation of sensory evidence leading up to a response. Neural activity recorded from lateral intraparietal cortex, frontal eye field and superior colliculus rises to a stereotyped threshold before response execution in non-human primates performing simple perceptual decision making tasks(Gold & Shadlen, 2007; Hanes & Schall, 1996; Huk, 2005; R. Ratcliff, 2003; Roitman & Shadlen, 2002; Schall, 2003). Similarly, electroencephalography (EEG) recordings from humans have found that the amplitude of the event-related centro-parietal-positivity (CPP), reaches a stereotyped threshold at the time of response, suggesting that the CPP tracks the accumulation of sensory evidence (Kelly & O’Connell, 2013; O’Connell, Dockree, & Kelly, 2012; Rungratsameetaweemana, Itthipuripat, Salazar, & Serences, 2018a). These experiments also suggest that the amount of available sensory evidence only influences the rate of evidence accumulation, not the response threshold (Kelly & O’Connell, 2013; Shadlen & Kiani, 2013), which is consistent with modeling results (Bogacz et al., 2006; Gold & Shadlen, 2007; Roitman & Shadlen, 2002; Shadlen & Kiani, 2013). Thus, these findings have been interpreted as empirical evidence that responses are triggered by the termination of evidence accumulation at a set threshold, irrespective of how quickly that threshold is reached. However, these experiments did not test whether this framework extends to decisions made under changing cognitive demands.

Indeed, simple modulations in response thresholds in a serial accumulate-to-threshold framework may not account for flexible decision making under speed pressure. Previous work using fMRI suggests that there are changes in both sensory and response processes with speed pressure (Forstmann et al., 2010; Ho et al., 2012; Ivanoff, Branning, & Marois, 2008; van Veen, Krug, & Carter, 2008), but assessing the timing of processes underlying decision making is difficult in fMRI studies as they do not have the temporal resolution to track evidence accumulation in real time. Importantly, one report manipulating the SAT performed single-unit recordings in the frontal-eye-field (FEF) of highly-trained non-human primates, a region containing a mixture of sensory, sensory-motor, and motor selective neurons (Heitz & Schall, 2012). Speed pressure increased baseline and maximum firing rates in most sensory neurons, and increased the magnitude and slope of firing rates at the time of response in motor selective cells encoding the impending saccadic response (Heitz & Schall, 2012). Note that these diverse results are inconsistent with accumulate-to-threshold models of the SAT, where speed pressure *decreases* response thresholds without modulating how sensory evidence accumulates (Heitz & Schall, 2012). However, the generalizability of this conclusion to other neural regions, varying levels of sensory evidence, and in subjects without extensive training is unknown. Additionally, previous studies have not dissociated decision making under external, explicit speed goals from internally regulated trial-to-trial variability in response times. Thus, it is unclear whether prior effects are related to changes in task demands or are instead due to distinct but correlated changes in general arousal or attentional state.

Here we manipulate speed pressure to test the assumptions of the traditional accumulate-to-threshold framework by measuring the CPP as a continuous index of evidence accumulation in human subjects. Along with speed pressure, we varied the amount of sensory evidence present on each trial to parametrically evaluate how speed goals impact the temporal sequencing of evidence accumulation and response execution. Finally, we isolate changes due to explicit speed goals from internally regulated variability by analyzing the data with and without matching response times across speed-emphasized and accuracy-emphasized conditions.

Our data suggest that when accuracy is emphasized, responses are made around the time when sensory evidence accumulation terminates. However, counter to accumulate-to-threshold models, responses are executed before evidence accumulation terminates under speed pressure. Importantly, response times are systematically shifted towards the time of maximal rate of evidence accumulation, indicating response processes maintain continual access to the state of evidence accumulation. Furthermore, decreasing the amount of available sensory evidence reduced response thresholds, not just accumulation rates, implying that reaching a stereotyped level of evidence does not automatically trigger a response. Together, these findings demonstrate a high degree of flexibility under changing speed goals inconsistent with serial frameworks of decision making.

## Results

### Contrast discrimination task

Twenty human subjects completed a visual contrast discrimination task while electroencephalography (EEG) signals were recorded (Figures 1A,B). Each trial started with a centrally presented full-field checkerboard stimulus flickering at 10Hz with 3.957 log (52% Michelson) contrast. After a pseudo-random interval of 600-1000 ms, the stimulus could increase, decrease, or maintain its contrast for an additional 2000 ms before stimulus offset. Subjects were instructed to discriminate the direction of a contrast change (two alternative forced choice: contrast increase or decrease) while maintaining fixation on a central fixation point.

**Figure 1:**
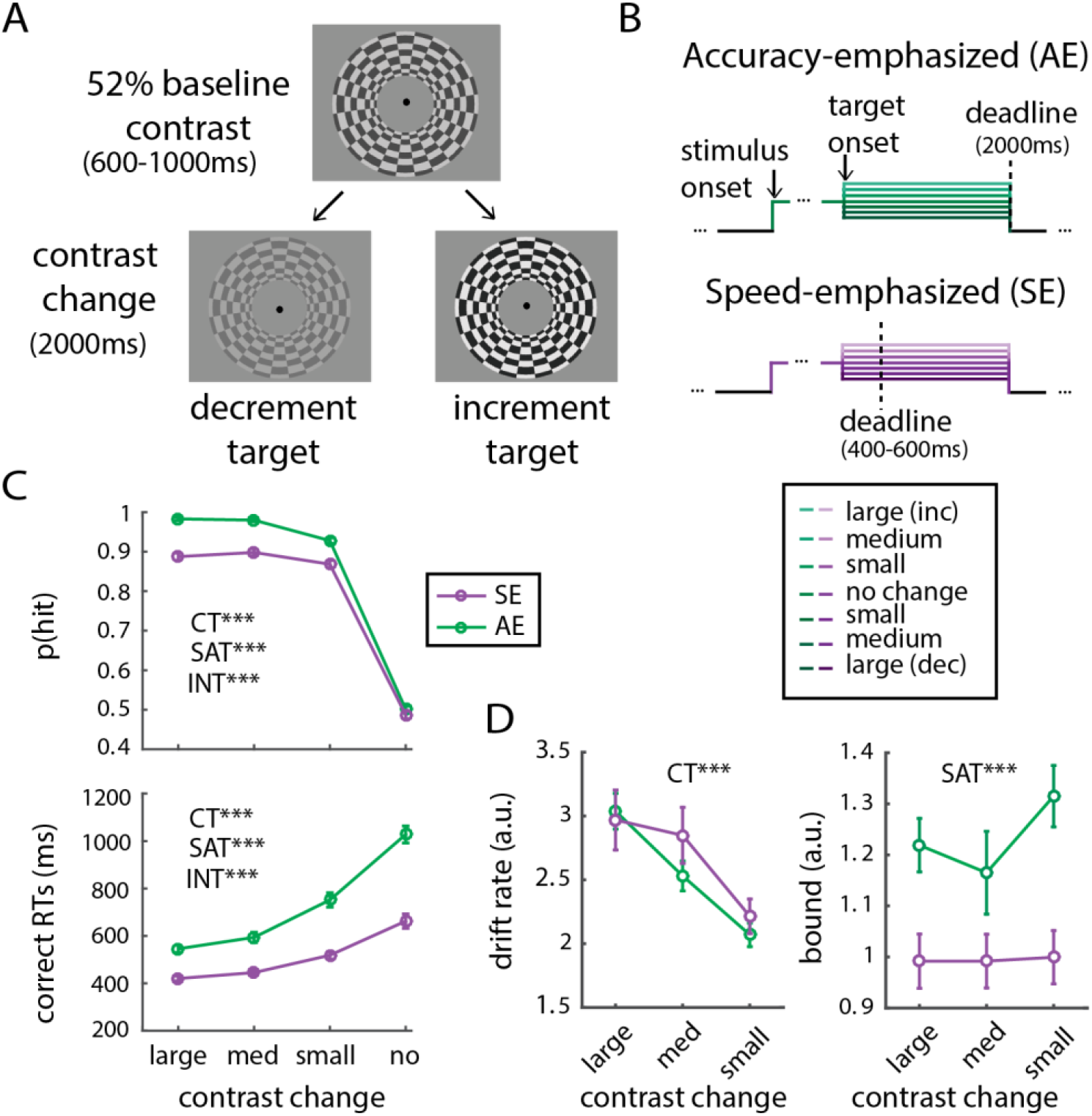
Task design, subject behavior, LBA modeling and SSVEP. Note: All error bars and shaded regions represent between subject standard error of the mean. **A**: Subjects completed a visual contrast discrimination task in which there was a contrast increment or decrement after a pseudo-random interval of 600-1000 ms from a 52% Michelson contrast baseline on each trial. This contrast increment or decrement could be small, medium, or large, and 1/7 of the trials were catch trials in which no contrast change occurred. **B:** Subjects were instructed to discriminate the direction of the contrast change (increment or decrement) while the 2000 ms the stimulus remained on the screen on accuracy emphasized blocks, or within 400-600 ms (jittered pseudo randomly) during speed emphasized blocks. Subjects were instructed that a correct discrimination within these time limits would lead to monetary reward. **C:** Here we plot hit rates (upper panel) and RTs (lower panel) collapsed across contrast increments and decrements (left panel). On no contrast change trials, hit rates were randomly assigned regardless of the subject’s response. Larger contrast changes lead to more accurate and faster responses. Additionally, subjects responded faster but less accurately with speed pressure (Purple line). Statistics were performed using a repeated measures ANOVA, and excluding no contrast change trials since the correct response is not well defined for these trials. CT = Contrast, SAT = Speed Accuracy Tradeoff and INT = interaction. **D:** Decreased sensory evidence leads to reduced drift rates while sensory evidence does not affect response threshold. Conversely, speed pressure reduces response threshold without impacting drift rates. Response threshold and drift rate parameters of the LBA model were fit using maximum likelihood estimation after collapsing behavioral data across increment and decrement trials within small, medium and large contrast change conditions. **C-D**: * indicates p<0.05, ** = p<0.01, *** = p<0.001 based on F statistics obtained with 2 way repeated measures ANOVA.

We explored how speed goals and amount of sensory evidence impact the temporal sequencing of evidence accumulation and response execution processes during perceptual decision making. Speed pressure was manipulated across experimental blocks. On speed emphasized blocks, subjects were instructed that correct responses would only be rewarded if made quickly after target onset (rewarded response interval was jittered pseudo-randomly from 400 to 600 ms after target presentation, following references (Heitz & Engle, 2007; Heitz & Schall, 2012)). During accuracy-emphasized blocks, subjects were instructed that responding correctly while the stimulus was on the screen would result in a monetary reward (i.e. within 2000 ms after target presentation). We varied the amount of sensory evidence across trials by showing either a small, medium, or large increase or decrease in the contrast of the stimulus. We presented each of the resulting seven levels of sensory evidence (including no change trials; see Methods) with equal probability. Note that while only correct trials are plotted for behavioral and EEG analyses, RTs from incorrect trials were used for modelling the behavioral data (see section below).

### Behavior

Behavioral data indicate that both our sensory evidence and SAT manipulations were successful (Figure 1C). There was a reliable effect of speed pressure on hit rates and RTs over all non-zero contrast change conditions (Figure 1C, Supplementary Figure 1A; post-hoc t-tests: t(19)’s = 2.92-10.89, all p’s ≤ 0.009 for hit rate; t(19)’s = 6.90-10.10, all p’s < 0.001 for RTs; two-tailed, FDR-corrected). As intended, the absolute, instead of signed, amount of sensory evidence was key in modulating behavior, indicated by consistent effects of speed goals and amount of sensory evidence across collapsed and non-collapsed behavioral data (evaluated with a repeated-measures ANOVA on SAT and contrast, reported below). Thus, for simplicity, we show data collapsed across contrast decrement and increment trials throughout the remainder of the text (unless otherwise noted).

As expected, larger changes in contrast led to faster and more accurate responses (Figure 1C, Supplementary Figure 1A; main effect of contrast: RT-F(2,38) = 137.78, p < 10^−15^, non-collapsed F(5, 95) = 63.18, p < 10^−15^: Hit Rate-F(2,38) = 25.64, p<10^−7^, non-collapsed F(5, 95) = 12.02, p < 10^−8^). Additionally, subjects responded faster but less accurately under speed pressure (main effect of SAT: RT - F(1, 38) =110.49, p<10^−8^, non-collapsed F(1, 95) = 107.16, p < 10^−8^; Hit Rate - F(1,38) = 189.67, p<10^−10^, non-collapsed F(1, 95) = 189.67, p <10^−10^). Finally, decreasing the amount of sensory evidence had a larger impact during accuracy emphasized blocks, leading to relatively lower hit rates and longer RTs (SAT x contrast interactions: Hit Rate - F(2,38) = 12.18, p<10^−4^, non-collapsed F(5, 95) = 14.34, p <10^−9^; RTs - F(2,38) = 40.53, p<10^−9^, non-collapsed F(5, 95) = 24.28, p < 10^−14^). We excluded no-change trials from these analyses to avoid spurious interactions since accuracy was randomly assigned with 50% probability on these trials (see Methods).

*Behavioral modelling suggests that speed pressure reduces response thresholds* To model how serial accumulate-to-threshold theories account for the impact of speed pressure and sensory evidence on behavior we estimated evidence accumulation rates (e.g. drift rates) and response thresholds using a Linear Ballistic Accumulator (LBA) model(Brown & Heathcote, 2008). The LBA specifies decisions as a race between independent accumulators corresponding to each possible choice in which the first accumulator to reach the response threshold determines decision choice and RT. We chose the LBA model due to its simplicity and ability to accommodate a variety of empirical results (Bogacz et al., 2010; Brown & Heathcote, 2008; Ho et al., 2012).

Importantly, the LBA model accounts for accuracy and RTs associated with both correct and incorrect responses, making it a natural choice for modeling responses reflecting the SAT(Brown & Heathcote, 2008). We fit each subject’s behavioral data using maximum likelihood estimation of response threshold and drift rate across SAT conditions and contrast change levels collapsed across increment/decrement trials (see ref(Donkin, Brown, & Heathcote, 2011), Methods).

We found that reductions in sensory evidence lead to slower drift rates (F(2, 38) = 33.03, p <0.001; Figure 1D) but had little impact on response thresholds (F(2, 38) = 2.67, p = 0.083). In contrast, speed pressure reduced response thresholds (F(1, 19) = 39.04, p <0.001) but did not influence drift rates (F(1, 19) = 0.73, p = 0.404). Thus, in line with existing computational models, our behavioral data can be accounted for by selective increases in drift rates with greater sensory evidence and reductions in response thresholds with speed pressure (Forstmann et al., 2010; Gold & Shadlen, 2007; Ho et al., 2012; O’Connell et al., 2012; Purcell et al., 2010; Rahnev & Denison, 2018; Rahnev, Evan, Riddle, Sue, & Esposito, 2016; Roger Ratcliff et al., 2016).

### Speed goals do not modulate early sensory responses

To evaluate whether speed goals and sensory evidence modulate early sensory encoding, we flickered the stimulus at 10Hz to measure the amplitude of the steady-state visually evoked potential (SSVEP; Figure 2A; see Methods; all FDR-correction from −500 to +2500 ms around the target). Consistent with past studies, SSVEP amplitude increased as a function of target contrast (timepoints from 66-481 ms and 830-2104 ms post-target survive FDR correction for the main effect of contrast: F(6, 114)’s = 2.46-11.57, p’s ≤ 0.028; (Di Russo, Spinelli, & Morrone, 2001; Sirawaj Itthipuripat, Garcia, Rungratsameetaweemana, Sprague, & Serences, 2014; Kim, Grabowecky, Paller, Muthu, & Suzuki, 2007; Norcia, Appelbaum, Ales, Cottereau, & Rossion, 2015; W. & J., 1972; Wang & Wade, 2011)). This indicates that increasing target contrast successfully increased the amount of sensory evidence represented in visual cortex. Importantly, speed pressure did not affect SSVEP amplitude (main effect of SAT: F(1, 19)’s ≤ 3.26, all p’s ≥ 0.087; SAT x contrast interaction: F(6, 114)’s ≤ 3.55, p’s ≥ 0.003, non-significant after FDR-correction, and note that uncorrected p’s ≤ 0.05 occurred long after the mean RT). This null result suggests that any effects of speed pressure on the CPP are not simply inherited from changes in early sensory gain.

**Figure 2:**
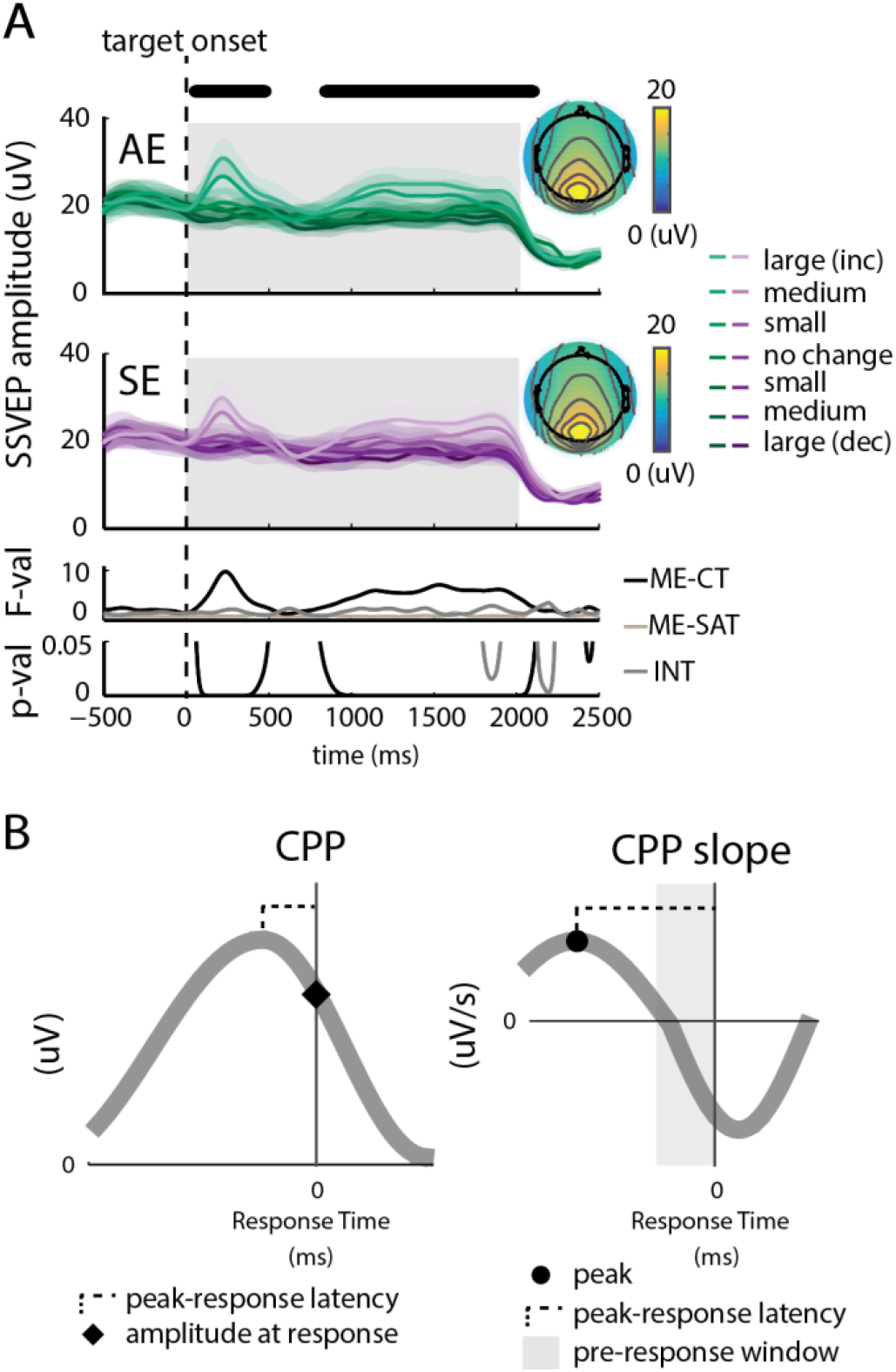
Early sensory evidence is not modulated by speed pressure and guide to interpreting CPP metrics **A:** We flickered the central stimulus at 10 Hz to measure SSVEP amplitude, an index of early sensory encoding. SSVEP amplitude from electrode Oz increased as a function of target contrast (indicated by black dots at top), while speed pressure did not significantly impact SSVEP amplitude after FDR correction. Bottom panels show F statistics and corresponding P values from a repeated measures 2 way ANOVA with contrast and SAT as factors. Inset topography plots show amplitude averaged over entire target duration (0 - 2000 ms, shaded gray). CT = Contrast, SAT = Speed Accuracy Tradeoff and INT = interaction, SE = Speed Emphasized, AE = accuracy emphasized. **B:** Guide to CPP Metrics. We show two idealized CPP examples in gray (left panel), with their corresponding slopes (right panel)

Although we did not find evidence that speed goals impact sensory gain, a previous study based on recordings in FEF of non-human primates, an area involved in sensory-motor integration as opposed to only sensory processing, did report an increase in sensory gain with speed pressure (Heitz & Schall, 2012; Schall, 2002; Thompson & Schall, 1999). To more closely emulate the analysis used in the previous study, we computed SSVEP amplitudes aligned to response onset (FDR corrected from −500 ms to +250 ms around response, see Methods). Similar to the stimulus-onset locked analysis presented above, higher contrast targets led to higher SSVEP amplitudes at the time of response onset (main effect of contrast: 2.44 ≤ F(6,114)’s ≤ 7.44, p’s ≤ 0.29 for timepoints from −457 to −17 ms pre-response that survive FDR correction). While there was also a modest pre-response interaction between contrast and SAT, this interaction did not survive FDR correction (Supplementary Figure 1B; F(6,114)’s ≤ 3.17, 0.007 ≤ p’s).

Even though this interaction did not survive FDR correction, we explored whether it could be due to changes in explicit task demands regarding speed pressure or to internal variability in attention or arousal. Therefore, we used a resampling procedure to match the RTs across speed and accuracy conditions to isolate effects due to changes in explicit task demands (see Methods, section: *RT sorting and RT matching analyses*). First, we observed a qualitative increase in SSVEP amplitude during trials with fast compared with slow RTs (Supplementary Figure 1C; Main effect of RT did not survive FDR correction: F(6,114)’s ≤ 9.56, 0.006 ≤ p’s; Contrast x RT interaction did not survive FDR correction: F(6,114)’s ≤ 3.73, 0.002 ≤ p’s). Moreover, when RTs were matched between speed and accuracy conditions, there was no longer any pre-response interaction between SAT and contrast (Supplementary Figure 1D; post-response interaction did not survive FDR correction: F(6,114)’s ≤ 2.94, 0.01 ≤ p’s). This suggests that the previous report of sensory gain in FEF neurons with speed pressure could be due in part to internally-regulated variability in RTs instead of explicit speed goals (Heitz & Schall, 2012).

*Sensory evidence, but not speed pressure, modulates CPP amplitude at response* Since our aim was to track the evidence accumulation process leading up to a response, we focus on the effects of speed goals and sensory evidence on the response-locked CPP (See Figure 2B for a guide to interpreting CPP metrics; Figures 3A-C; we present detailed target-locked CPP results in the Supplementary Results). First, we found that CPP amplitude at response onset, posited as an empirical measure of response threshold, decreased with the magnitude of target contrast (Figures 3C,D; F(3, 19) = 96.67; p <10^−15^). This putative modulation of response thresholds is inconsistent with accounts that sensory evidence selectively modulates the speed of evidence accumulation (Drugowitsch et al., 2012; Kelly & O’Connell, 2013; R. Kiani et al., 2008; O’Connell et al., 2012). Note that accumulate-to-threshold models sometimes incorporate response thresholds that collapse as deliberation time increases, which may account for our data given that we found longer RTs on trials with low contrast targets (Drugowitsch et al., 2012; Roger Ratcliff et al., 2016). However, reduced CPP amplitude with lower sensory evidence is still observed in RT-matched data, indicating that collapsing response thresholds with long RTs cannot account for this effect (F(3, 57) = 92.37; p <10^−15^; Figure 3F, Supplementary Figure 2A,D;). Also contrary to accumulate-to-threshold models of the SAT, speed pressure did not significantly reduce CPP amplitude at the time of response (Figure 3D; F(1,57) = 3.31, p = 0.085; SAT x contrast interaction: F(3, 57) =1.44, p = 0.241; also note gaps in significance at response time in Figures 3A-C). Interestingly, post-hoc t-tests suggest a slight increase in the amount of sensory evidence needed to trigger a response instead of the decrease suggested by computational models (e.g. the LBA in present study; (Hanks, Kiani, & Shadlen, 2014;

**Figure 3:**
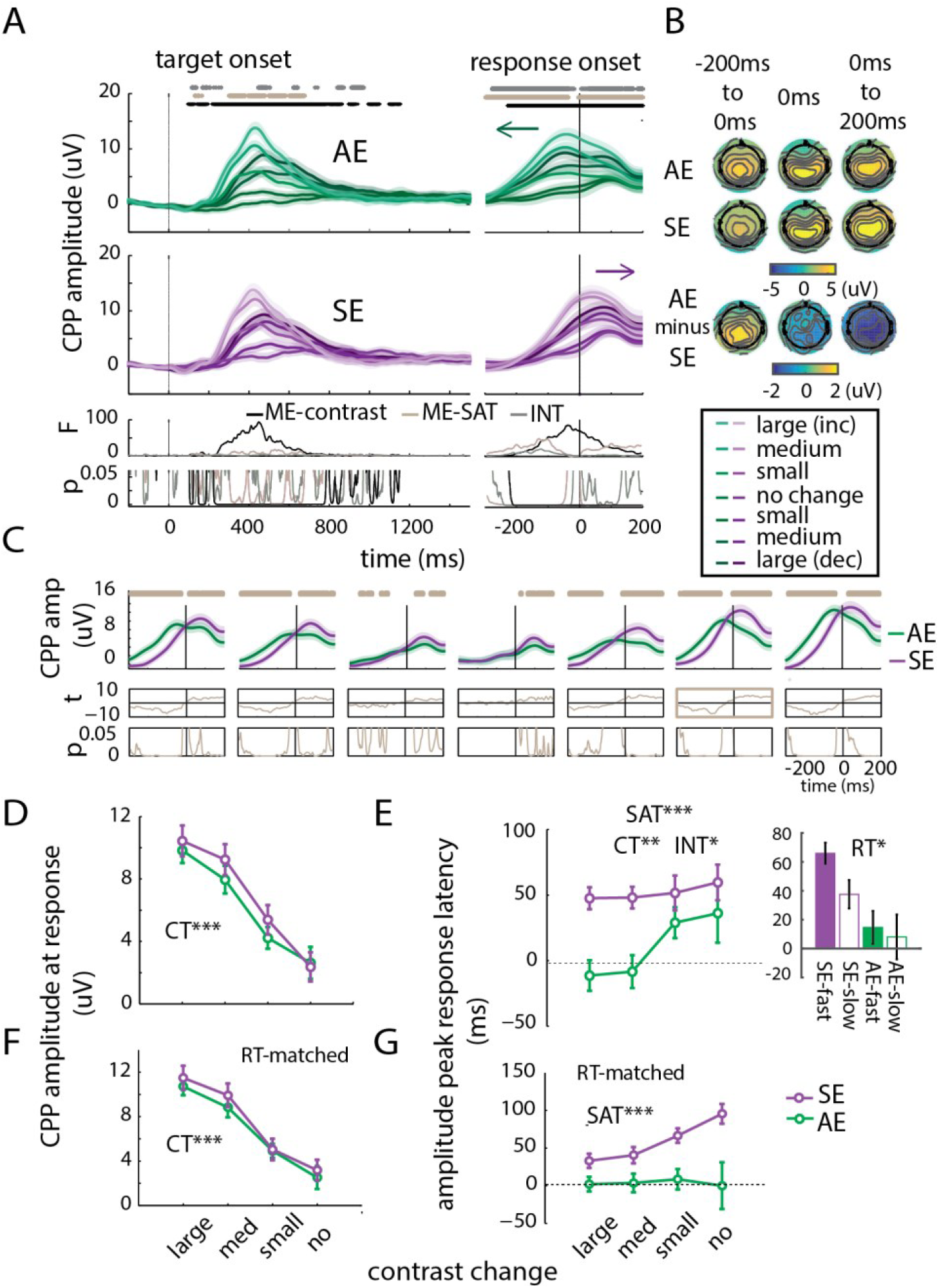
Speed pressure shifts response locked Central Parietal Positivity. Note: All error bars and shaded regions represent between subject standard error of the mean. **A:** Target locked (left panel) and response locked (right panel) CPPs split by accuracy emphasized (AE, top panel) and speed emphasized (SE, middle panel) and contrast level. Bottom panel shows F statistics and corresponding P values for a repeated measures 2-way ANOVA with contrast and SAT as factors, dots above top panel indicate significance after FDR correction. **B:** Response locked CPP amplitude topographic maps separated by accuracy emphasized (top panel), speed emphasized (middle panel) conditions averaged over all contrast conditions in 3 temporal epochs: from −200 ms to response onset, at response onset, or averaged from response onset to 200 ms after response onset. Topography for accuracy minus speed emphasized is plotted in the bottom panel. **C:** Response-locked CPP amplitude for each SAT condition plotted separately for each contrast level (subplots are ordered from largest contrast decrement to largest contrast increment, with no change trials in the center; top panel). T statistic and corresponding p-value are plotted in the bottom panels, dots above panels indicate significance after FDR correction. Vertical line indicates response time. **D:** CPP amplitude at response onset decreases with sensory evidence, but not speed pressure. Amplitude at response onset shown here are averaged within contrast magnitudes and across increment and decrement conditions. **E:** Response time was subtracted from the time of CPP peak and averaged within contrast magnitudes-note that negative peak-to-response latencies indicate that peaks occurred before the response, and vice versa. Peak-to-response latencies for CPP amplitude are closer to zero with high levels of sensory evidence when accuracy is emphasized (left panel). Trials were median split by RT within SAT condition and collapsed over contrast conditions. We found graded peak-to-response latencies such that faster responses exhibited longer peak-to-response latencies (right panel). **F:** CPP amplitudes at response were averaged as in D, but here only trials in which RTs were equal for speed and accuracy conditions were considered (see methods). **G:** CPP peak-to-response latencies as in E, but here only trials in which RTs were equal for speed and accuracy conditions were considered. With matched RTs, peak-to-response latencies during accuracy emphasized blocks are not significantly different from zero regardless of contrast level. **D-G**: * indicates p<0.05, ** = p<0.01, *** = p<0.001 based on F statistics obtained with 2 way repeated measures ANOVA.

R. Ratcliff, 2003)), consistent with previous evidence in non-human primates (Heitz & Schall, 2012) (t(19)’s = −1.09, −2.49, −2.15, p = 0.29, 0.02, 0.04 for large, medium and small contrast changes). That said, there was no significant effect of speed pressure on CPP amplitude at response over all contrast levels (F(1,57) = 3.31, p = 0.085; SAT x contrast interaction: F(3, 57) =1.44, p = 0.241), which held when we matched RTs between speed and accuracy conditions (Figure 3F, Supplementary Figure 2A,B; F(1, 19) = 1.9, p = 0.184). Importantly, note that CPPs actually continued to rise after the responses were made under speed pressure, a result inconsistent with the traditional accumulate-to-threshold theory in which responses are only triggered once sensory accumulation reaches its peak (Figure 3A-C).

### Speed pressure shifts response timing relative to the CPP

Instead of modul0061ting CPP amplitude at the time of a response, speed pressure appears to temporally shift the CPP relative to response (Figures 3A-C,E). While responses occurred close to the CPP peak in the accuracy condition, speed pressure shifted response times approximately 52 ± 11 ms before the CPP peak, quantified as peak-to-response latency (mean ± SE; Figure 3E left; SAT main effect on amplitude peak-to-response latency: F(1, 19) = 14.34, p = 0.001). After median splitting trials into fast and slow bins we found a graded pattern such that explicitly speeded trials with fast RTs exhibited the largest shifts in peak-to-response latencies (Figure 3E right; Main effect of RT: F(1, 19) = 7.63, p = 0.012; see Methods, section: *RT sorting and RT matching analyses*). Importantly, any nonzero peak-to-response latencies during accuracy emphasized trials were eliminated after matching RTs, while the shift in peak-to-response latency with speed pressure was more prominent – increasing to 59 ± 13 ms (Figure 3G, Supplementary Figure 2A; Main effect of SAT: F(1, 19) = 19.08, p <0.001; t-tests against zero-accuracy: t(19)’s = 0.20, 0.33, 0.94 and 0.36 with p’s = 0.20, =0.33, =0.94 and = 0.36 for large, medium, small and no contrast change, respectively; speed: t(19)’s = 3.62, 3.73, 6.94 and 7.34 with p’s < 0.002, <0.002, <10^−5^, <10^−6^ for large, medium, small and no contrast change). This robust effect after matching RTs indicates this shift cannot be accounted for by confidence or urgency signals that are known to be associated with RTs (Baranski & Petrusic, 1994; Drugowitsch et al., 2012; Roozbeh Kiani, Corthell, & Shadlen, 2014). Together, these results suggest that subjects can flexibly shift response timing with respect to the ongoing accumulation process under explicit speed pressure to respond before evidence is fully accumulated (Cosman, Arita, Ianni, & Woodman, 2016).

### Sensory evidence does not robustly affect the timing of responses relative to the CPP

Reducing the amount of sensory evidence also shifted responses to occur before the peak of the CPP (F(3, 57) = 4.12, p = 0.010, SAT x contrast interaction: (F(3, 57) = 2.85, p = 0.046). However, unlike the shift associated with speed pressure, the effect of sensory evidence is attenuated when RTs are matched across conditions, suggesting that the shift was largely attributable to trials with slow RTs on low contrast accuracy emphasized trials (Figure 3E,G, Supplementary Figure 2A; F(3, 57) = 3.65, p = 0.018, SAT X contrast interaction: F(3,57) = 1.87, p > 0.14).

### Both sensory evidence and speed pressure modulate the slope of the CPP

We next explored the slope of the CPP, a proxy for the rate of evidence accumulation, as a potential mechanism by which speed pressure flexibly modulates the timing of responses with respect to the ongoing evidence accumulation process. We numerically computed the slope of the response-locked CPP (Figure 4A; see Methods) and found that mean CPP slope preceding responses was higher with both speed pressure and increased sensory evidence (averaged from −100-0 ms leading up to the response, Figure 4B; F(1, 19) = 50.70, p < 10^−6^; F(3, 57) = 18.47, p <10^−7^, (Gold & Shadlen, 2002, 2003, 2007; Heitz & Schall, 2012; Purcell et al., 2010; Roitman & Shadlen, 2002; Shadlen et al., 2006)). Note that while there was an interaction between speed pressure and sensory evidence on pre-response CPP slope (F(3, 57) = 18.47, p < 10^−9^), post-hoc t-tests revealed this was because speed pressure increased CPP slope in all contrast conditions except for the “no change” condition (t(19)’s = −6.37, −7.55, −4.85 and −1.39 with p’s < 10^−5^, <10^−6^, <0.001 and = 0.18 for large, medium, small and no contrast change, respectively). Importantly, the increase in pre-response CPP slope held even when we matched RTs between speed and accuracy conditions (Figure 4E, Supplementary Figure 2E; SAT: F(1, 19) = 51.08, p < 10^−6^; Contrast: F(3, 57) = 14.5, p <10^−6^). Again, post-hoc t-tests revealed significant effects of speed pressure in all contrast change conditions except for the no-change trials (t(19)’s = 5.82, 7.84, 3.52, and 1.60 with p’s < 0.0001, <10^−6^, = 0.002, = 0.127 for large, medium, small, and no contrast changes, respectively). This increase in pre-response CPP slope indicates that explicit speed pressure increases the rate of evidence accumulation just before a response, an effect that could be due to differences in either the magnitude of the peak slope or the timing of the peak slope relative to response.

**Figure 4:**
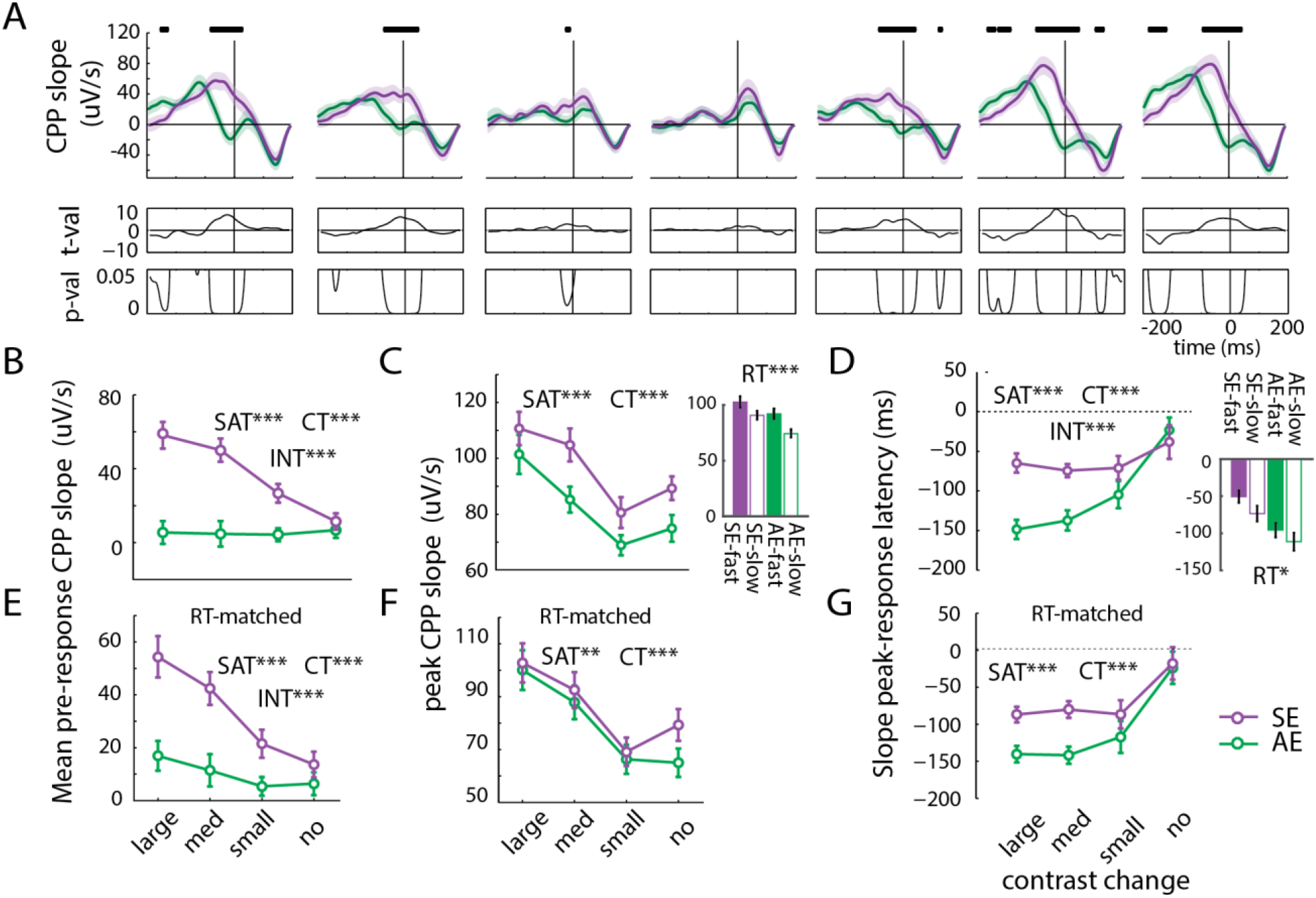
Both sensory evidence and speed pressure modulate CPP slope. Note: All error bars and shaded regions represent between subject standard error of the mean. **A:** We numerically approximated the derivative of the response-locked CPP. Timepoints for which slope on speed and accuracy trials are significantly different according to a paired t-test are plotted with dots above the top panel, while corresponding t-statistics and p-values are plotted in middle and bottom panels (subplots are ordered from largest contrast decrement to largest contrast increment, with no change trials in the center). **B:** Mean CPP slope leading up to a response increased with both speed pressure and sensory evidence. **C:** Peak CPP slopes, irrespective of timing, are qualitatively distinct from mean pre-response drift (left panel). This is quantitatively shown in a lack of an interaction effect. Trials were median split by RT within SAT condition and collapsed over contrast conditions, revealing that faster RTs also increase peak CPP slope. **D:** Speed pressure shifts peak-to-response latencies closer to the time of peak CPP slope, and this effect is more pronounced on large and medium contrast change trials (left panel). Furthermore, median splitting trials by RT within SAT condition and collapsed over contrast conditions revealed that faster RTs also bring peak-to-response latencies closer to zero (right panel). **E:** As in B, CPP slope leading up to a response increased with both speed pressure and sensory evidence, but here only Speed and Accuracy emphasized trials with equal RTs were considered. **F:** As in B, CPP slope leading up to a response increased with both speed pressure and sensory evidence, but here RTs were equated across speed and accuracy emphasized trials. However, note that these effects were not robust within contrast conditions, as can be seen qualitatively (t(19)’s = 0.66, 1.35, 0.42, and 1.89, p’s = 0.519, 0.191, 0.679, 0.074 for large, medium, small and no contrast change, respectively). **G:** As in D, speed pressure and low contrast shifts peak-to-response latencies closer to the time of peak CPP slope. **B-G**: * indicates p<0.05, ** = p<0.01, *** = p<0.001 based on F statistics obtained with 2 way repeated measures ANOVA.

### Explicit speed pressure does not increase the magnitude of the peak CPP slope

To better understand how speed pressure increases mean pre-response CPP slope, we first consider the magnitude of the peak CPP slope, which increased with both speed pressure and greater sensory evidence (Figure 3C left; SAT: F(1, 19) = 15.55, p < 0.001; CT: F(3, 57) = 20.88, p < 10^−8^; no interaction F(3, 57) = 0.86, p =0.46). However, note that the modulatory pattern of speed goals on the maximal slope did not show the interaction with contrast seen previously in the pre-response slope (lower contrasts reduced peak slope during both SAT conditions, while contrast only impacted mean pre-response slope with speed emphasis -Figure 4B vs 4C). Thus, increases in pre-response CPP slope due to speed pressure cannot be solely attributed to increases in the magnitude of the peak slope. Indeed, matching RTs greatly reduced the link between the maximal CPP slope and speed pressure such that even though there was still a significant effect of speed pressure on the magnitude of the peak CPP slope (F(1, 19) = 8.52, p =0.009), these results were not robust within each contrast condition (Figure 4F, Supplementary Figure 2E; post-hoc t-tests: t(19)’s = 0.66, 1.35, 0.42, and 1.89, p’s = 0.519, 0.191, 0.679, 0.074 for large, medium, small and no contrast change, respectively). Moreover, when trials were split into fast and slow RTs, we found that fast responses increase the magnitude of the peak CPP slope independent of speed pressure (Figure 4C right; main effect of RT: F(1,19) = 45.13, p < 10^−5^, RT x SAT interaction: F(1,19) = 1.6, p = 0.2; see Methods Section: *RT sorting and RT matching analyses*). This significant increase in the magnitude of peak CPP slope with fast RTs suggests internal processes like attention and arousal may increase the maximal rate of evidence accumulation. Additionally, matching RTs reveals that explicit speed pressure doesn’t robustly modulate the magnitude of peak CPP slope, meaning increases in maximal accumulation rate could be a selective mechanism underlying trial-to-trial variability during perceptual decision making.

### External speed pressure shifts responses toward the time of peak CPP slope

As the magnitude of peak CPP slope was not robustly impacted by explicit speed goals, we next explore the timing of peak CPP slope as a potential source of the increase in pre-response CPP slope with explicit speed pressure. There is a qualitatively apparent shift in the timing of responses with respect to CPP slope under speed goals (Figure 4A). Indeed, speed pressure systematically shifts responses 40.7 ± 9.1 ms closer to the time of peak CPP slope compared with accuracy emphasized trials (Mean ± SE; Figure 4D; Main effect of SAT: F(1, 19) = 27.32, p < 0.0001). Additionally, the shift in RT with speed pressure was more pronounced on trials with higher amounts of sensory evidence (Main effect of Contrast: F(3, 57) = 11.7, p < 10^−5^; SAT x Contrast interaction: (F(3, 57) = 7.98, p < 0.001; Figure 4A,D left; t(19)’s = −6.14, −5.07, −2.02 and −0.80, with p’s <10^−5^, <0.001, = 0.058 and = 0.44 for large, medium, small and no contrast changes, respectively). As before, we dissociate external, explicit speed goals from internally regulated variability in RT by re-computing the timing of the peak CPP slopes after matching RTs (Supplementary Figure 2E). Highly consistent with the non RT matched data, we found that speed pressure brought response times closer to the time of peak CPP slope (F(1, 19) = 17.22, p < 0.001), an effect driven by trials with more sensory evidence (t(19)’s = 3.78, 5.64, 1.29, and 0.20 with p’s = 0.001, <0.0001, = 0.212, = 0.844 for large, medium, small, and no contrast changes, respectively). Splitting trials into fast and slow RTs revealed that internally regulated fast responses only shifted responses 19 ms ± 4.3 SE closer to the time of peak slope as compared with slow trials (Figures 4C-D right; main effect of RT: F(1,19) = 7.67, p = 0.012). Thus, there seem to be distinct mechanisms through which humans make speeded responses: internally regulated fast decisions increase the rapidity of evidence accumulation, while responses execution is shifted to occur once evidence accumulation decelerates under external, explicit speed pressure.

## Discussion

The present results suggest that, when under speed pressure, responses can be executed before evidence accumulation is complete instead of being automatically triggered by its termination. This is evident in a shift of the CPP relative to responses such that responses are made shortly after evidence accumulation reaches its maximal rate. This pattern diverges from predictions made by computational models within the accumulate-to-threshold framework, including our own LBA model of behavioral data, which explain faster responses made under explicit speed pressure by a simple reduction in the amount of accumulated sensory evidence. Instead, our observation that evidence continues to accumulate even after speeded responses are made argues in favor of an architecture in which humans can flexibly engage in response planning concurrently with evidence accumulation to meet task demands.

Interestingly, our analyses of the amplitude and slope of the CPP indicate that explicit speed goals modulated response timing in a systematic way. With speed pressure, subjects responded once evidence accumulation began to decelerate, even though sensory evidence was still accumulating. Alternatively, when accuracy was emphasized responses were not executed until sensory evidence accumulation was complete. This shifting of response times is a potential temporal mechanism by which speed pressure decreases reliance on incoming sensory evidence (Ho et al., 2012; Ivanoff et al., 2008) to execute responses as soon as the rate of sensory evidence accumulation starts to decelerate. Our observation that additional sensory information is discounted with speed pressure makes sense because accumulating evidence takes time, suggesting decisions are made once a point of “diminishing returns” is reached. Further experiments are needed to determine whether the criterion for diminishing returns is always anchored to the rate of evidence accumulation, as we found with explicit speed pressure, or whether other factors are involved under various task demands.

We note the possibility that the observed shifts with speed pressure could reflect changes in decision urgency or confidence if the CPP reflects a decision signal that isn’t selective for sensory evidence accumulation (Donchin, Kubovy, Kutas, Johnson, & Tterning, 1973). However, responses made under speed pressure occurred even farther before the peak of the CPP instead of being eliminated when RTs were matched between speed and accuracy conditions, a factor known to be strongly correlated with urgency and confidence. Additionally, when accuracy was emphasized the data are largely consistent with previous characterizations of the CPP as an abstract decision signal that reaches a maximal level at the time of behavioral response, suggesting that speed pressure alters the timing between evidence accumulation and response processes in a way that diverges from accumulate-to-threshold modelling frameworks.

While further work is needed to determine the neural generators of the CPP and its functional significance, the present data suggest that, at least in the context of this task, serial accumulate-to-threshold theories do not capture the flexibility of decision making under changing cognitive demands.

It is important to note that while both internally regulated variability in RT and explicit, externally imposed speed goals affect the slope of the CPP, they do so in dissociable ways. Explicit speed pressure shift response timing with respect to the ongoing evidence accumulation process. On the other hand, internal processes leading to fast RTs independent of externally imposed speed goals appear to result from more rapid sensory evidence accumulation (Kelly & O’Connell, 2013). This indicates a need for future studies to dissociate explicit speed goals due to task demands from internally mediated variability in RTs.

We also found that the total amount of sensory evidence accumulated, and not just the speed of evidence accumulation, decreased monotonically with less available sensory evidence largely independent of reaction times. This suggests that the amount of evidence humans accumulate is proportional to the total amount of sensory evidence available in the environment, and further experiments are needed to elucidate the mechanism by which this accumulation threshold is set. Notably though, this finding contrasts with predictions from traditional accumulate-to-threshold models in which changing the amount of sensory evidence impacts the rate of evidence accumulation but leaves response thresholds unchanged.

Together, our results suggest an alternative to the accumulate-to-threshold models that dominate current perceptual decision making theory. Instead, we suggest that downstream decision processes operate in parallel to the accumulation of sensory evidence, allowing for flexible sampling of evidence accumulation to adjust response timing according to task demands.

## Methods

### Subjects

21 human adults, who had normal or corrected-to-normal color vision, were recruited to participate in the experiment. All subjects provided written informed consent as required by the local Institutional Review Board at the University of California, San Diego (UCSD; IRB#110176), and the experiment was conducted under protocols following the Declaration of Helsinki. Subjects were compensated $15 per hour for participation plus an additional monetary reward of up to $15 depending on behavioral performance (see details below). The data of one subject was not included in the analysis because the subject decided to terminate their participation before the experiment was completed. This left 20 subjects in the final analyses (8 males; 2 left-handed; mean age = 22.62 years old, SD = 6.47).

### Stimuli and tasks

We presented stimuli and tasks on a PC running Windows XP using MATLAB (Mathworks Inc., Natick, MA) and the Psychophysics Toolbox (version 3.0.8; Brainard 1997; Pelli 1985). Subjects were seated 60 cm from the CRT monitor (which had a grey background of 34.51 cd/m^2^, 120Hz refresh rate) in a sound-attenuated and electromagnetically shielded room (ETS Lindgren).

Subjects performed a contrast discrimination task where they had to report the direction of a contrast change (either decreasing or increasing contrast) of a centrally presented checkerboard stimulus. This checkerboard stimulus was flickered at 10 Hz using equal on-off duty cycles of 6 frames each. The target always appeared at the on-cycle of the SSVEP flicker frequency, and lasted for 2000ms. Each trial began with a black circle fixation at the center of the computer screen where they were instructed to fixate for 600, 700, 800, 900 or 1000 ms (randomly drawn on each trial). After the fixation period, a checkerboard stimulus of 3.957 log contrast (52% Michelson) contrast appeared for 600–1000 ms (i.e., a non-target period) followed by either a small contrast decrement/increment of 0.181 log contrast (rendered at 44%/63% Michelson contrast), medium contrast decrement/increment of 0.362 (37%/75% Michelson), or large decrement/increment of 0.543 (31%/90% Michelson), or the same non-target stimulus rendered at the same contrast (no change). On each trial, a particular target contrast (including no change) appeared with an equal probability of 1/7.

SAT was manipulated across blocks of 70 trials. In speed emphasized blocks, Subjects had up to 400-600ms (jittered pseudo-randomly) after target onset to respond to gain monetary reward. In accuracy emphasized blocks, subjects had 2000ms after target onset (i.e. until target offset) to respond to gain monetary reward. Any given correct response made before the deadline yielded a monetary reward of 0.01 cents, leading to an average reward of $9.73 ± 0.72 (mean ± SD).

### EEG preprocessing

After data collection, data from the scalp electrodes were re-referenced to the algebraic mean of the two mastoid electrodes. Then, the raw time series from each electrode was bandpass filtered between 0.25 to 58 Hz to attenuate slow drift and 60 Hz line noise. After filtering, data were epoched into 5 second intervals from −1.5 to +3.5 seconds around target onset. Exclusion criteria for trials were based on thresholds determined for each subject by visual inspection. Trials were excluded from ERP analyses if the EOG channels located above or below either eye reached on average 111.5 ± 23.2 mV (mean ± SD for blinks) or if the EOG channels located outside either outer canthi reached 76.25 ± 12.86 mV (mean ± SD for saccades). Using these criteria, an average of 15.3% ± 10.5% S.D. of trials were excluded, leaving 1162.5 ± 150 (mean ± SD) trials for EEG analyses.

### EEG analysis

SSVEP amplitudes were estimated from data averaged over trials within each contrast and SAT condition at central occipital electrode Oz using a wavelet transform with a Gaussian frequency domain kernel with 2 Hz FWHM (0.8493 Hz SD)(Di Russo et al., 2001; Sirawaj Itthipuripat et al., 2014; Kim et al., 2007; Norcia et al., 2015; W. & J., 1972; Wang & Wade, 2011). The absolute values of these complex coefficients were then taken to obtain SSVEP amplitude measures for each contrast and SAT condition. For RT locked SSVEPs, data were averaged after realigning each trial to the first on-frame of each duty cycle nearest to the RT on that trial.

Event-related-potentials (ERPs) were computed by taking the average time courses centered on either the target time (target-locked) or the behavioral response (response-locked). For the response locked data, we re-epoched each trial from −300 ms to +200 ms around the behavioral response on that trial. Averages were taken within each contrast and SAT condition, leading to an average of 71.9 ± 13.1 SD trials per each SAT x Contrast condition without sorting by RTs, and 35.9 ± 6.5 SD after sorting based on median RT. For plotting purposes, data were filtered using a stop pass third order Butterworth filters between 8 and 12 Hz to attenuate the SSVEP signal, and a low pass third order Butterworth filter at 22 Hz after averaging. However, reported statistics were computed on unfiltered ERPs. Here, we used the CPP component as a neural index of sensory evidence accumulation following previous studies(Kelly & O’Connell, 2013; O’Connell et al., 2012; Rungratsameetaweemana, Itthipuripat, Salazar, & Serences, 2018b; Twomey, Murphy, Kelly, & O’Connell, 2015). The CPP data were analyzed at centro-parietal electrode (CPz) based on previously published studies(S. Itthipuripat, Ester, Deering, & Serences, 2014; S Itthipuripat, Cha, Rangsipat, & Serences, 2015; Kelly & O’Connell, 2013; O’Connell et al., 2012). Note that a decision-variable with similar accumulation dynamics to those seen in the CPP has also been observed in a late positive deflection or P3 component(Twomey et al., 2015), which has a long history in the ERP literature as being related to decision signals(Cosman et al., 2016; Donchin et al., 1973; Friedman, Simson, Ritter, & Rapin, 1975; Sirawaj Itthipuripat, Cha, Byers, & Serences, 2017; Sirawaj Itthipuripat, Cha, Deering, Salazar, & Serences, 2018; G. R. Mangun & Hillyard, 1988; George R. Mangun & Buck, 1998; Nelli, Itthipuripat, Srinivasan, & Serences, 2017; Squires, Donchin, & Squires, 1977).

RT sorted data were obtained within-subject by splitting trials into fast and slow bins according to each subject’s median RT determined separately for each contrast and SAT condition. We first combined data from the speed and accuracy emphasized trials and then divided all trials based on a median split of RTs at each contrast level. We then collapsed across all contrast levels, after performing the median split, to avoid confounding RT and contrast.

RT matched data were obtained for each subject by computing a vector ranging from the minimum to maximum RTs across speed and accuracy conditions in 100 ms steps, leading to an average of 6.71 ± 1.45 SD bins. Then, the number of speed trials and the number of accuracy trials were computed for each bin, and the minimum number of trials was taken. For each RT bin and SAT condition, that minimum number of trials was then randomly selected, without replacement, from the entire set of available RTs for each condition, and this process was repeated 100 times and then averaged across all the resampling iterations. Finally, trials from all RT bins were averaged within contrast and SAT condition (39.11 ± 9.42 trials, average ± SD).

CPP amplitude at RT was extracted separately for each contrast and SAT condition, and then averaged over large, medium and small contrast increment and decrement change trials separately. Similarly, peak CPP amplitude and its latency were estimated separately for each contrast condition and then averaged over contrast increment and decrement trials.

CPP slope was approximated by using the built-in ‘diff’ function in Matlab. This yeilded an estimate of the CPP derivative at each sample t by subtracting its amplitude at sample t from amplitude at sample t+1. We then multiplied the estimated derivative by the EEG sampling rate to convert units from mV/sample to mV/second.

### Statistics

F statistics were computed using a 2 way repeated measures ANOVA with a combination of SAT (2 levels), contrast conditions (4 or 7 levels), or RT (2 levels) as independent variables as reported in the text. T statistics were reported using paired t-tests. When these statistics were computed over multiple timepoints we used FDR correction at an alpha level of 0.05 to control for multiple comparisons. P values are reported as < 10^−15^ when below the machine precision. In all figures, error bars and shaded regions represent between subject standard error of the mean.

## Acknowledgements

We thank Kai-Yu Chang for help with data collection, and Margaret Henderson, Angus Chapman and Timothy Sheehan for feedback on the manuscript. Supported by NEI R01-EY025872 to J.T.S and a James S. McDonnell Foundation award to J.T.S, by an HHMI International Fellowship and the Royal Thai Scholarship from the Ministry of Science and Technology in Thailand to S.I, and by NSDEG graduate fellowship to S.N.

## Supplementary Materials

### Supplementary Results

#### Target Locked Cento-parietal positivity (CPP) event-related potential

We observed a robust CPP component that emerged after target onset and was maximal at central posterior electrode Pz (Figure 2A left, Figure 2C, Supplementary Figure 3A Middle row). As done with response-locked results, we collapsed across contrast decrement and increment trials to simplify statistical analyses since the effects of sensory evidence and speed goals were similar. We report results according to the magnitude of the contrast change (large, medium and small changes).

We first report the peak amplitude for CPPs locked to the time of target onset. We found a decrease in peak CPP amplitude and an increase in target-to-peak amplitude latency with decreasing magnitude of contrast change (Supplementary Figure 3B; F(3, 57) = 48.81, p < 10^−12^; Supplementary Figure 3D, F(3, 57) = 69.51, p < 10^−15^). Speed pressure significantly increased peak CPP amplitude (Supplementary Figure 3B; F(1, 19) = 21.25, p <0.001), but had no effect on target-to-peak latency (Supplementary Figure 3D; F(1, 19) = 4.08, p = 0.06).

Note that these contrast and SAT – dependent modulations in CPP could be precisely predicted by the peak of the normalized RT distributions (Supplementary Figure 3A Top Row). Specifically, the peak of the RT distribution decreased with increasing contrast change (Supplementary Figure 3C; F(3,57) = 318.09, p < 10^−15^) and decreased with speed pressure (F(1, 19) = 28.49, p < 10^−4^). Also, consistent with the result seen with CPP target-to-peak latency, the peak latency of the RT distributions increased as the degree of contrast change decreased Supplementary Figure 3E; F(3, 57) = 22.66, p < 10^−9^) and we found a robust effect of speed goals on the peak latency of the normalized RT distributions (F(1, 19) = 36.82, p < 10^−5^).

To control for this relationship between response execution and peak CPP amplitude, we matched mean RTs between the speed and accuracy conditions. We still found a decrease in peak CPP amplitude and an increase in target-to-peak amplitude latency with decreasing magnitude of contrast change (Supplementary Figure 3F, F(3, 57) = 59.57; Supplementary Figure 3G, p < 10^−15^, F(3, 57) = 18.45, p < 10^−7^). However, there was no main effect of SAT on CPP peak amplitude or its latency (F(1, 19) = 3.66, p = 0.071, F(1, 19) = 0.39, p = 0.54). This suggests that the increase in CPP amplitude and the marginal decrease in CPP latency with speed pressure was primarily driven by a narrower RT distribution (first row in Supplementary Figure 3A).

**Supplementary Figure 1:**
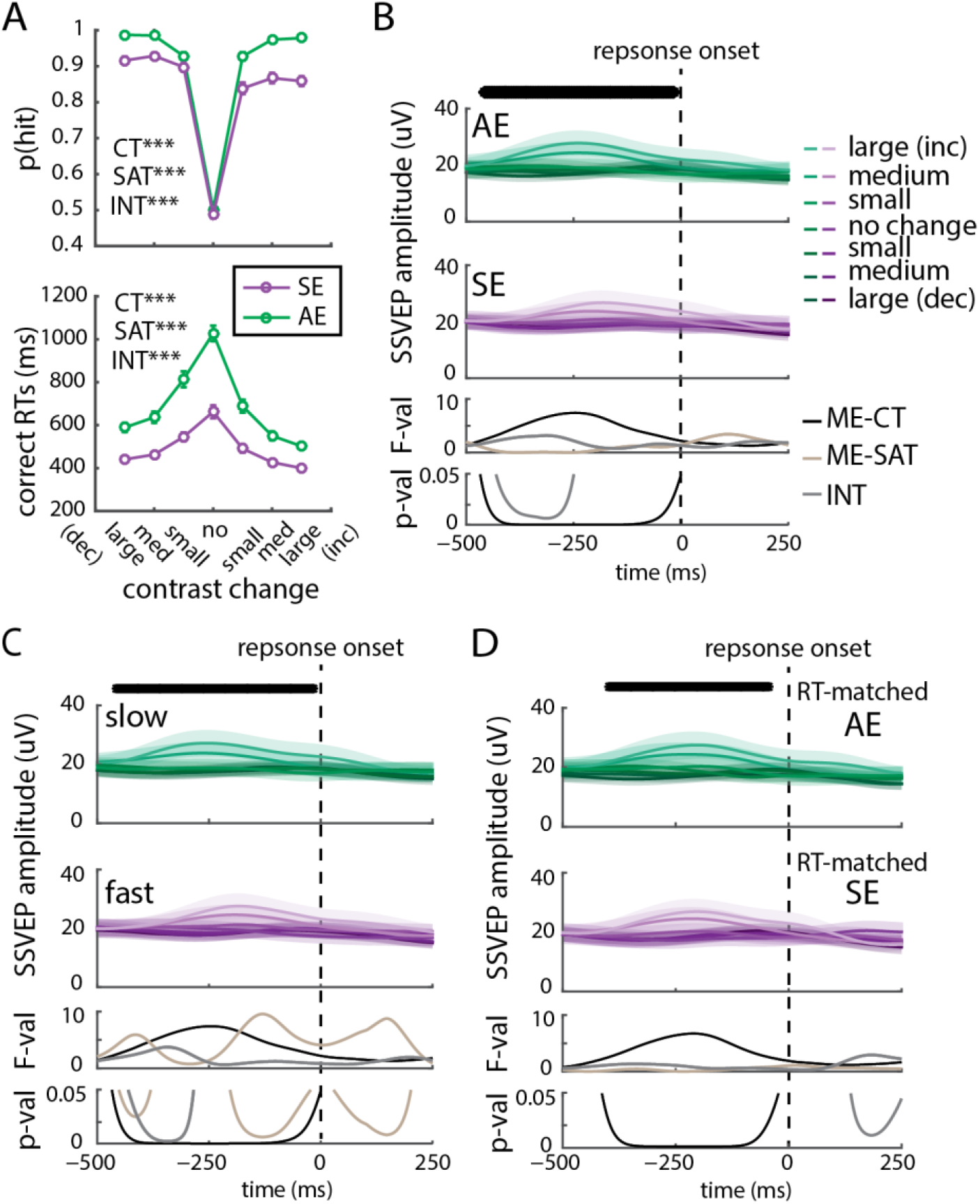
Behavior and RT-locked early sensory evidence. **A:** Here we plot hit rates (upper panel) and RTs (lower panel) collapsed across contrast separately for each of the seven contrast conditions (right panel). On no contrast change trials, hit rates were randomly assigned regardless of the subject’s response. Larger contrast changes lead to more accurate and faster responses. Additionally, subjects responded faster but less accurately with speed pressure (Purple line). Statistics were performed using a repeated measures ANOVA, and excluding no contrast change trials since the correct response is not well defined for these trials. B: SSVEP amplitude was recomputed after re-aligning each trial to the first frame of each cycle nearest to the RT on that trial. SSVEP amplitude from electrode Oz increased as a function of target contrast (indicated by black dots at top), while speed pressure did not significantly impact SSVEP amplitude after FDR correction. Bottom panels show F statistics and corresponding P values from a repeated measures 2 way ANVOA with contrast and SAT as factors. **C:** After splitting trials by RT, SSVEP amplitude from electrode Oz increased as a function of target contrast (indicated by black dots at top), while RT did not significantly impact SSVEP amplitude after FDR correction. Bottom panels show F statistics and corresponding P values from a repeated measures 2 way ANVOA with contrast and RT as factors. **D:** After matching RTs, SSVEP amplitude from electrode Oz increased as a function of target contrast (indicated by black dots at top), while speed pressure did not significantly impact SSVEP amplitude after FDR correction. Bottom panels show F statistics and corresponding P values from a repeated-measures 2 way ANVOA with contrast and SAT as factors.

**Supplementary Figure 2:**
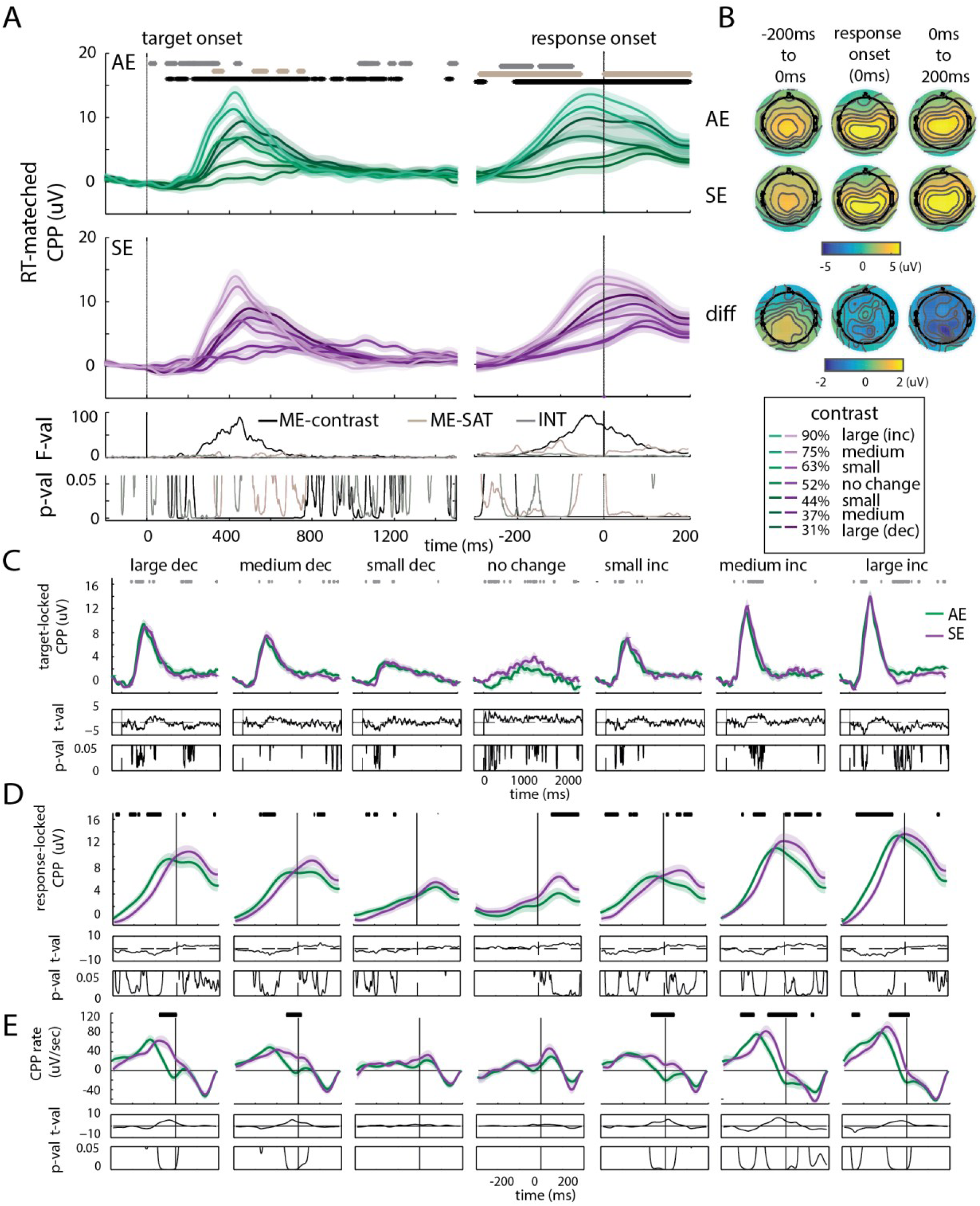
RT-matched CPP results. Note: All error bars and shaded regions represent between subject standard error of the mean. **A:** RT matched target locked (left panel) and response locked (right panel) CPPs split by accuracy emphasized (AE, top panel) and speed emphasized (SE, middle panel) and contrast level. Bottom panel shows F statistics and corresponding P values for a repeated measures 2-way ANOVA with contrast and SAT as factors, dots above top panel indicate significance after FDR correction. **B:** RT matched response locked CPP amplitude topographic maps separated by accuracy emphasized (top panel), speed emphasized (middle panel) conditions averaged over all contrast conditions in 3 temporal epochs: from −200 ms to response onset, at response onset, or averaged from response onset to 200 ms after response onset. Topography for accuracy minus speed emphasized is plotted in the bottom panel. **C:** RT-matched target-locked CPP amplitude for each SAT condition plotted separately for each contrast level (top panel). T statistic and corresponding p-value are plotted in the bottom panels, dots above panels indicate significance after FDR correction. **D:** RT-matched response-locked CPP amplitude for each SAT condition plotted separately for each contrast level (top panel). T statistic and corresponding p-value are plotted in the bottom panels, dots above panels indicate significance after FDR correction. **E:** RT-matched response-locked CPP slope each SAT condition plotted separately for each contrast level (top panel). T statistic and corresponding p-value are plotted in the bottom panels, dots above panels indicate significance after FDR correction. **C-E:** Subplots are ordered from largest contrast decrement to largest contrast increment, with no change trials in the center.

**Supplementary Figure 3:**
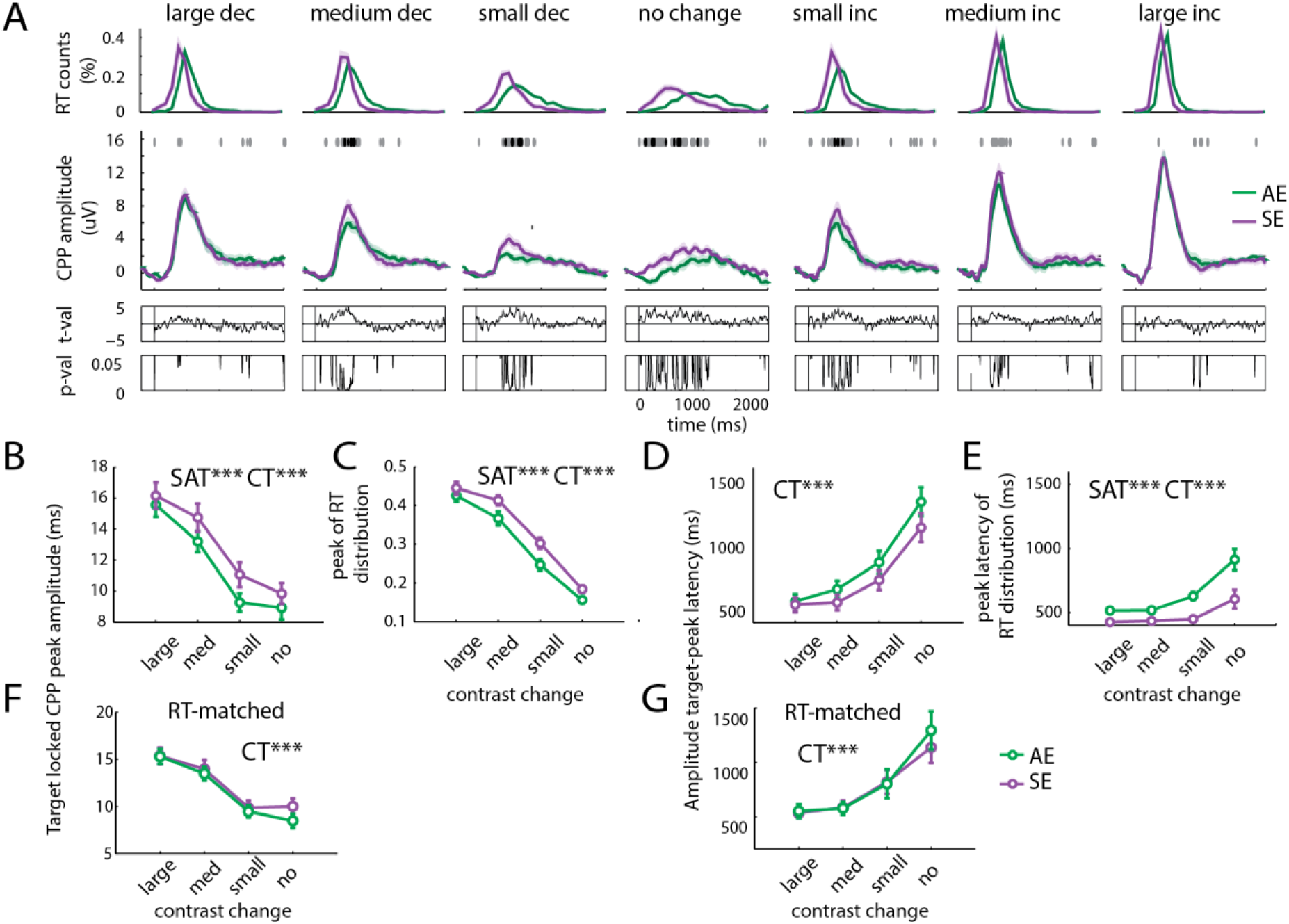
Target locked CPP metrics. Note: All error bars and shaded regions represent between subject standard error of the mean. **A:** RT distributions separated by speed and accuracy emphasis conditions and contrast change level. Distributions were normalized such that the y-axis represents the percentage of responses made at that time within condition (top panel). Target-locked CPP timecourses for each SAT condition plotted separately for each contrast level (middle panel). T statistic and corresponding p-value are plotted in the bottom panels, dots above panels indicate significance after FDR correction (Bottom Panels). RT matched target locked CPP amplitude. Subplots are ordered from largest contrast decrement to largest contrast increment, with no change trials in the center. **B:** Target-locked CPP peak amplitude increases with both sensory evidence and speed pressure. Amplitudes here are averaged within contrast magnitudes and across increment and decrement conditions. **C:** As shown for CPP amplitude in B, the peak magnitude or concentration of the RT distribution also increases with both sensory evidence and speed pressure. **D:** The latency from target to CPP peak amplitude increases with contrast but is not affected by speed pressure. **E:** The latency, or value, of the peak of the RT distribution decreases with both contrast and speed pressure. **F:** Target-locked CPP peak amplitude as in B, but using only trials with matched RTs between speed and accuracy conditions. While there is still an increase of amplitude with sensory evidence, there is no longer an effect of speed pressure on peak amplitude. **G:** Target-locked CPP peak latency as in D, but using only trials with matched RTs between speed and accuracy conditions. **B-G**: * indicates p<0.05, ** = p<0.01, *** = p<0.001 based on F statistics obtained with 2 way repeated measures ANOVA.

**Author Contributions** SN analyzed the data and wrote the manuscript. SI conceived and implemented the experiment, collected the data, analyzed the data, and co-wrote the manuscript. NR analyzed the data and co-wrote the manuscript. JTS conceived and supervised the project and co-wrote the manuscript

